# Coupling lysosomal polarity and purinergic signaling regulates B cell activation

**DOI:** 10.64898/2026.09.15.751411

**Authors:** M. Alamo Rollandi, J.P. Bozo, I. Riobó, T. López-López, J.M. Kux, M.Y. Jaeckstein, B. Rissiek, J. Heeren, D. Sauma, P.J. Sáez, M.I. Yuseff

## Abstract

B cell activation is initiated by engagement of the B cell receptor (BCR) with immobilized antigens, which triggers changes in cell polarity promoting the establishment of an immune synapse that is further shaped by signals from the microenvironment. However, how cell polarity coordinates the sensing of extracellular cues to regulate B cell activation remains unclear. Here, we investigated the impact of adenosine triphosphate (ATP), a conventional danger signal related to inflammation, on B cell function. We found that ATP released by B cells, as well as exogenously added ATP, acts as a promoter of the extraction and presentation of immobilized antigens. Endogenous ATP was produced by mitochondria recruited at the immune synapse and locally released via Pannexin 1 channels to sustain purinergic signaling at the immune synapse. We identified P2RX4 trafficking to the plasma membrane through the Rab6a+ trans-Golgi network and VAMP7+/LAMP1+ lysosomes as a key step in this response. In addition, the local activation of P2RX4 at the immune synapse triggered a migratory switch from motile to sessile, suggesting that this receptor acts as a negative regulator of B cell migration. These findings reveal ATP as a local enhancer of B cell function, and an unexpected role of P2RX4 as a molecular switch for B cell activation.

## Introduction

Cell polarity is the asymmetric distribution of intracellular components, which organizes signaling, forces, and a plethora of cellular responses including cell division, communication, migration^1,2^. Thus, organelle polarity and function determine cellular behavior in health and disease, including epithelial function, the immune response, and cancer metastasis^3^. In lymphocytes, cell polarity plays a major role during antigen presentation, and the local release of degradative enzymes at the immune synapse is required for proper antigen extraction, degradation, and presentation^4,5^.

B lymphocytes are key components of adaptive immunity and require efficient cell polarity for antibody secretion and T cell activation^6^. Their activation begins when the B cell receptor (BCR) recognizes antigen, triggering immune synapse formation, antigen internalization and processing, and the subsequent presentation of antigen-derived peptides on MHC II molecules to cognate T cells ^7,8^. The microenvironment can shape B cell interactions with antigens and immune synapse formation. For example, mechanical cues, such as substrate stiffness^9,10^, and extracellular matrix components like galectin-8^11^. can modulate B cell activation, antigen extraction, and MHC-II presentation. Innate signals further shape these responses, as TLR activation can synergize with BCR signaling^12^, whereas TLR9 activation can impair antigen capture and presentation, limiting CD4+ T cell interactions and humoral immunity^13,14^.

A key inflammatory mediator released during tissue damage is ATP, which in its extracellular form at high concentrations acts as a damage-associated molecular pattern (DAMP), triggering purinergic signaling pathways^15^. Through activation of purinergic (P2) receptors, ATP regulates several cellular processes, including chemotaxis of innate immune cells, such as neutrophils, dendritic cells, and macrophages. Additionally, ATP signaling through P2 receptors regulates T cell and macrophage activation and function, promoting the secretion of inflammatory interleukins such as IL-1β by supporting inflammasome assembly and maturation^16–18^.

Despite extensive characterization of purinergic signaling in innate immune cells and T cells, its role in B cell biology remains largely unexplored. Studies examining the direct effects of ATP on B cell activation have yielded conflicting results^19,20^. Nevertheless, indirect evidence suggests that ATP could regulate key steps involved in antigen extraction; ATP has been shown to promote cytoskeletal remodeling through P2X7-dependent signaling and downstream kinase activation^21,22^ as well as to regulate lysosomal dynamics^23^. By modulating these processes, ATP may also influence antigen extraction and presentation in B cells. Although B cells express multiple P2 receptors^24^, it remains unclear whether extracellular purinergic cues directly modulate their activation and antigen-presenting capacity.

Here we identify extracellular ATP as both an autocrine signal triggered by lymphocyte activation, and an enhancer of this response once acting as a danger signal. B lymphocyte activation required P2RX4 function and specific subcellular localization. P2RX4 localized at the lysosomes or plasma membrane determined a functional switch from a motile to a sessile state, which were associated with antigen processing and presentation, respectively. This ATP-P2RX4-dependent switch was required for efficient lymphocyte activation and antigen presentation and emerges as a key regulator of immune cell function.

## RESULTS

### Extracellular ATP promotes antigen extraction and presentation in B cells

B cells acquire antigens from the surface of APCs in secondary lymphoid organs, where extracellular ATP is released by APCs or generated as a result of tissue damage and inflammation^17,25,26^. Although ATP is present in the B cell microenvironment, its effect on their cellular functions and activation remains unclear. To understand its role on B cell activation, we first evaluated whether B cells locally release ATP.

For this purpose, we used a mouse B cell line (IIA1.6)^27,28^, and latex beads coated with a BCR+ ligand (IgG), which is a well-established in vitro system to mimic the formation of an IS. BCR-ligand (IgM), was used as a negative control. These cells were transfected with a sensor for extracellular ATP, which increases fluorescence upon extracellular ATP binding^29^. Using live-cell imaging, we found a significant increase in the sensor’s fluorescence intensity at the synapses of activated cells, indicating that ATP is being released locally at the antigen-contact site (Fig. 1a,b). Similar results were obtained when B cells were activated on antigen-coated coverslips, in which ATP also accumulated at the synapse under activating conditions (Extended Data Fig. 1a,b,c).

**Figure 1:**
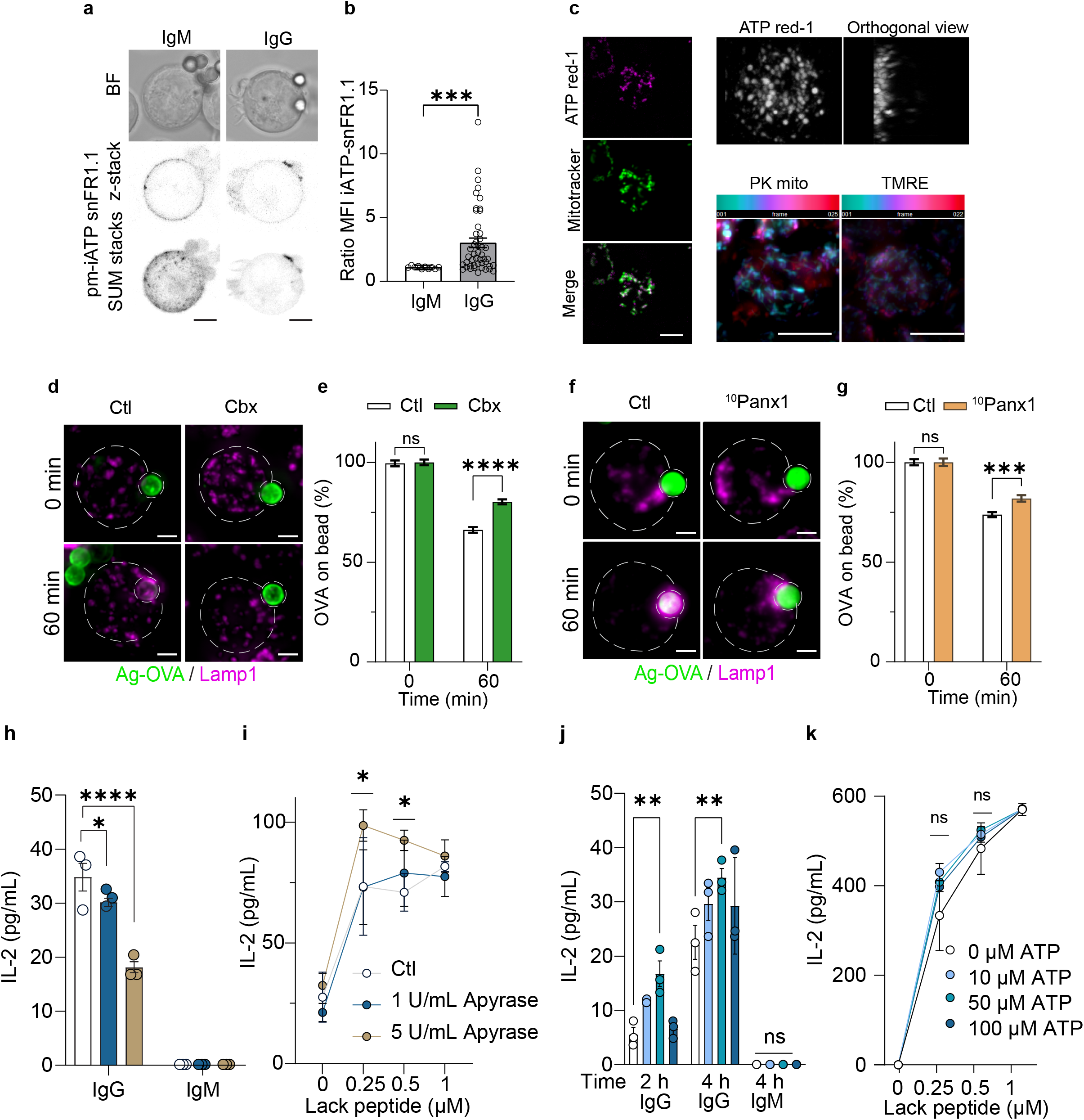
Extracellular ATP promotes antigen extraction and presentation in B cells. (**a**) Representative confocal images of B cells expressing the ATP sensor (pm-iATPsnFR1.1) activated with IgG or IgM coated beads for 60 min. Scale bar 3 μm. (**b**) Fold change of the mean fluorescence mean intensity (MFI) of the ATP sensor within the 3 μm diameter of the coated bead interacting with cells. t-test. ***P <0.001. Error bars are mean ± SEM. (**c**) Representative widefield live acquisition of B cells activated on IgG-coated slides. Left panel: IS plane visualization of cells stained with ATPred-1 and mitotracker. Right panel: IS plane and orthogonal view of cells stained with ATPred-1 (top). Z-plane color code for cells stained with PK mito and TMRE (bottom). (**d-g**) Shows the effect of Cx and Panx blockers over antigen extraction. Representative widefield images of B cells in the presence of 10 µM of Cbx (a non-selective Cx and Panx blocker) and (**f**) 200 µM of ^10^Panx1 (a Panx1 mimetic peptide) activated with IgG-OVA-coated beads for the indicated times. OVA (green) and Lamp1 (magenta). (**e**) and (**g**) Quantification of OVA fluorescence intensity remaining on beads interacting with cells. n ≥ 20 cells pooled from N = 3 independent experiments. Scale bar 3μm. Two-way ANOVA with Sidak’s multiple comparison test. P values illustrated with asterisks are ** <0.01, *** <0.001, and **** <0.0001. Error bars are mean ± SEM. (**h**) Antigen-Lack presentation assay of B-cells in the presence of 1 and 5 U/mL of apyrase and (**j**) with 10, 50 and 100 µM ATP. B-cells were incubated with IgG-Lack-coated beads for 2 and 4 h. Cells were fixed and co-incubated with LMR 7.5, Lack-specific T-cells, for 4 h at 37 °C. (**i**)and (**k**) Respective Lack peptide assay for the conditions shown. IL-2 levels were measured by ELISA. Mean amounts of IL-2 are shown as representative of three independent experiments performed in triplicate. Two-way ANOVA with Sidak’s multiple comparison test.

To explore the potential sources for the local release of ATP by B cells, we focused on mitochondria, as the main source of ATP^30^. First, we co-stained mitochondria with mitotracker and ATPred-1, a fluorescent dye that labels ATP^31^. In B cells activated on antigen-coated coverslips for 60 min, we observed mitochondria recruited to the IS plane (Fig. 1c left), which colocalized with the ATPred-1 dye. Orthogonal views of the ATPred-1 depict ATP-rich mitochondria polarized towards the IS, as shown independently using both morphology-based enhanced mitochondrial visualization and membrane potential–dependent mitochondrial markers, PKmito and TMRE (Fig. 1c right). This suggests that metabolically active mitochondria are recruited to the IS and may contribute to the local ATP pool released by B cells.

Next, we evaluated possible channels involved in mediating ATP release by B cells. Pannexin (Panx1) and Connexin (Cx) forming-channels contribute to immune cell activation, and Cx43 and Panx1 are also expressed in B cells and mediate their activation^32–35^. Thus, we evaluated whether these channels contributed to the ability of B cells to extract immobilized antigens. B cells were treated with carbenoxolone (Cbx), which inhibits both Cx and Panx channels, lanthanum (III) (La^3+^), a broader blocker of hemichannels, or with the Panx1-selective blocker ^10^Panx1 (Fig. 1d,e,f,g and Extended Data Fig. 1d,e,f). Cells were then stimulated with beads containing OVA and a BCR-specific ligand (antigen-OVA-coated), in the presence of the blockers. Antigen extraction efficiency was then evaluated by measuring the residual fluorescence of OVA on the beads following their interaction with treated B cells. Pharmacological blockade with either Cbx, La^3+^ or ^10^Panx1, significantly reduced the ability of B cells to extract immobilized antigen (Fig. 1d,e,f,g and Extended Data Fig. 1d,e,f), suggesting that channel-mediated ATP release by B cells contributes to their activation.

To assess whether ATP released by B cells impacts the extraction of immobilized antigens, B cells were treated with apyrase, an ATP-hydrolyzing enzyme. To ensure efficient depletion of ATP, B cells were preincubated with 1 or 5 U/mL of apyrase for 10 min before stimulation with antigen-OVA-coated beads. Treatment with apyrase (5 U/mL) significantly reduced the ability of B cells to extract immobilized antigen at 60- and 120-min following activation compared to control cells (Extended Data Fig. 1g,h). We next evaluated whether the effect of apyrase translates into reduced antigen presentation on MHC-II molecules to CD4+ T cells and performed an antigen presentation assay, as previously described^36^. B cells were stimulated with IgG- or IgM-Lack-coated beads. After 3 h of stimulation with the beads in the presence of 1 and 5 U/mL of apyrase, cells were fixed and co-incubated with the CD4+ T cell line LMR7.5, which recognizes a Lack-derived peptide presented on MHC-II. IL-2 levels in the supernatant, reflecting T cell activation, were measured by ELISA. We found that B cells exposed to high concentrations of apyrase significantly reduced their antigen-presentation capacity to T cells, with 5 U/mL of apyrase showing higher peptide presentation than control or 1 U/mL apyrase treatment, indicating that this defect did not result from diminished surface expression of MHC-II. Instead, it stemmed from reduced antigen extraction and processing by B cells (Fig. 1h,i). Together, these results suggest that upon BCR activation, B cells release ATP, which potentiates their capacity to extract and present antigens.

Having established that B cells release ATP into the extracellular space, a process associated with enhanced antigen extraction and impaired by inhibition of ATP-release channels, we next asked whether exogenous ATP could reproduce these effects. We therefore examined how extracellular ATP, derived from non-B cell sources, could influence antigen extraction by B cells. To this end, cells were pre-treated for 10 min with increasing concentrations of ATP (10 to 100 µM) before stimulation with antigen-OVA-coated beads for different times (0, 30, 60, 120 min). This range is sufficient to mimic ATP levels released from neighboring cells and reflects conditions observed during acute inflammation^37^. Antigen extraction was quantified by measuring the fluorescence of residual OVA on the beads following interaction with B cells. We found that stimulation with 10 µM of ATP increased antigen extraction throughout all time points (Extended Data Fig. 1i,j). The extraction of immobilized antigens by B cells depends on the recruitment of lysosomes towards the antigen contact site and their fusion at the synaptic membrane^36^. Thus, we next evaluated whether this increase in antigen extraction was due to increased lysosomal docking at this region. As anticipated, ATP increased lysosome accumulation at the IS after 60 min, relative to control cells (Extended Data Fig. 1i,l).

Because ATP is rapidly metabolized by ectonucleotidases (CD39/ENTPD1 and CD73/NT5E), we next assessed their expression in B cells. Gene expression from the ImmGen database revealed that most leukocytes predominantly express CD39 over CD73 (Extended Data Fig. 1l). We confirmed expression by flow cytometry analysis of resting B cells, which revealed minimal surface expression of either enzyme (≤ ∼2% CD39^+^ or CD73^+^) (Extended Data Fig. 1m left). In contrast, intracellular staining showed that ∼0% and ∼90% of cells expressed CD73 and CD39, respectively (Extended Data Fig. 1m middle). Additionally, we found that 30% of CD73 is located at the surface of activated B cells (Extended Data Fig. 1m right). To study the effects of extracellular ATP independently of its hydrolysis, we used a slowly hydrolyzable analogue of ATP, adenosine 5′-O-(3-thiotriphosphate) (ATPγS), and evaluated its impact on antigen extraction. We found that stimulation with ATPγS enhanced antigen extraction at both early (30 min) and late (120 min) stages of activation (Extended Data Fig. 1n). Notably, this effect was not accompanied by increased lysosome accumulation at the antigen contact site (Extended Data Fig. 1o), suggesting that ATPγS affects functional quality rather than quantity of lysosomes recruited to the synaptic interface as shown by Cathepsin activity assay using the substrate Magic Red. (Extended Data Fig. 1p,q). To validate our findings under more physiological conditions, we examined the effect of ATPγS in primary B cells isolated from the spleens of wild-type BALB/c mice. In BALB/c B cells, treatment with 10 µM ATPγS displayed enhanced antigen extraction at early time points (15 min) of B cell activation, whereas higher concentrations (50 and 100 µM), increased extraction at later time points of activation (30 and 120 min) (Extended Data Fig. 1r,s,t). Overall, primary B cells recapitulate the behavior observed in the cell line, supporting a role for extracellular ATP in increasing antigen extraction.

To determine whether extracellular ATP enhances antigen presentation to CD4⁺ T cells, B cells were stimulated with IgG- or IgM-Lack-coated beads, as described before. After 2-4 h of stimulation in the presence of increasing concentrations of ATP (10–100 µM), B cells were fixed and co-incubated with the CD4+ T-cell line. IL-2 levels in the supernatant, reflecting T cell activation, show that ATP significantly enhanced antigen presentation, particularly at 50 µM, with no discernible impact of ATP on peptide presentation, indicating that this enhancement was not due to increased MHC II surface expression but rather reflects improved antigen extraction and processing by B cells (Fig. 1j,k).

Altogether, these results show that ATP enhances the ability of B cells to extract and present immobilized antigens. Notably, this effect is mediated, at least in part, by ATP being released by B cells through ATP-permeable hemichannels, as inhibition of these channels significantly impairs antigen extraction. The P2RX4 receptor is required for antigen extraction and presentation.

Having shown that extracellular ATP enhances the capacity of B cells to extract and present immobilized antigens, we next sought to identify the receptor that is mediating this response. So, we focused on the purine-sensitive P2X receptor family, which are ionotropic and mainly activated by ATP, but with different affinities^37^. Using the ImmGen database, a gene expression heatmap revealed that most immune cells predominantly express P2RX4 and P2RX7, with mature B cells highly expressing P2RX4 (Fig 2a and Extended Data Fig. 2a).

**Figure 2:**
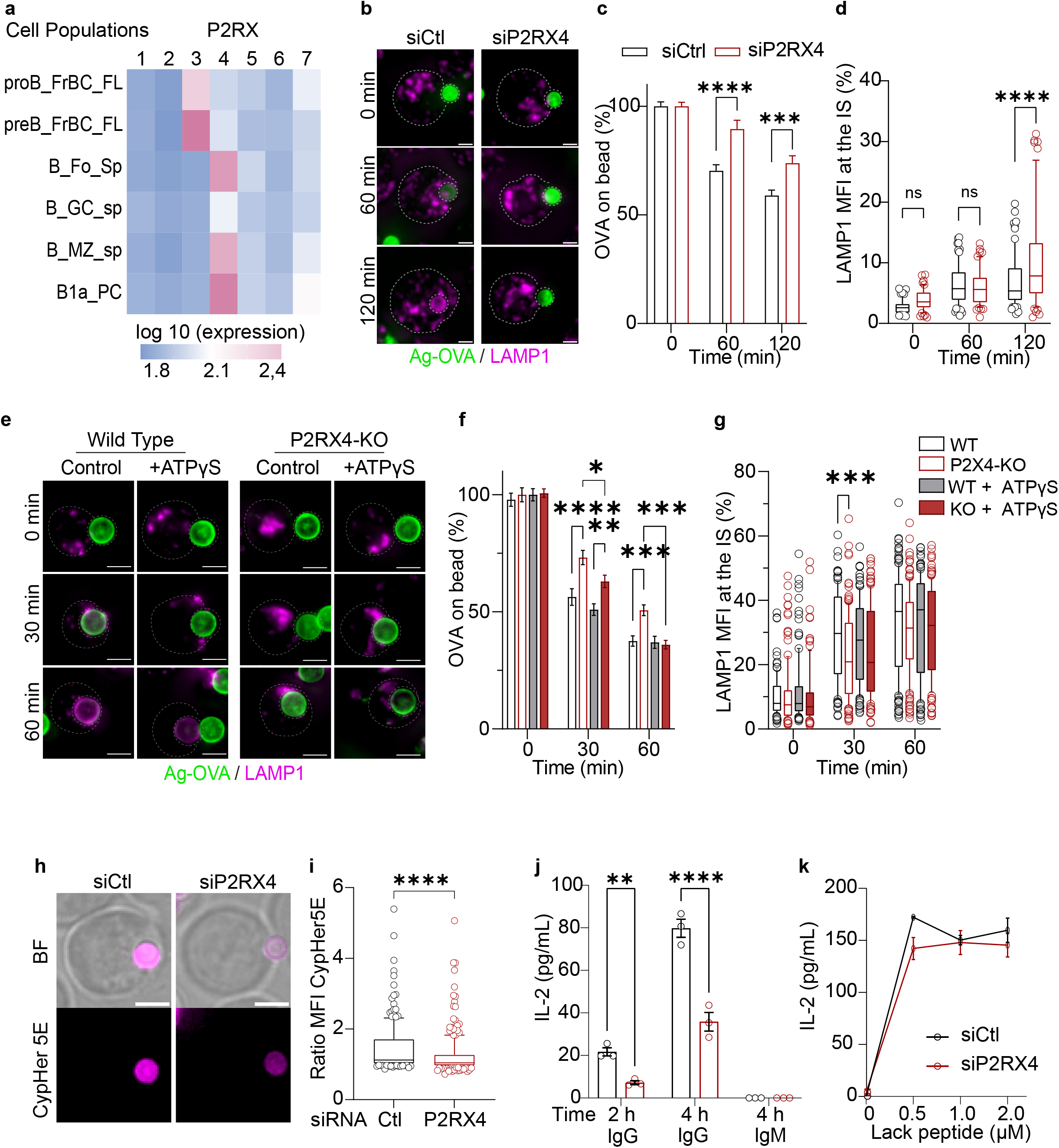
The P2RX4 receptor is required for antigen extraction and presentation. (**a**) Gene heatmap expression of P2X receptor family in B cell populations. (**b**) Representative widefield images of B cells silenced for P2RX4 activated with IgG-OVA-coated beads for the indicated times. OVA (green) and Lamp1 (magenta). (**c**) Quantification of OVA fluorescence intensity remaining on beads interacting with cells. (**d**) Quantification of the percentage of the MFI of Lamp1 in the area enclosed by the bead normalized by total MFI of Lamp1 in the cell. (**e**) Representative wide field images of primary B cells isolated from the spleen of wild-type and P2RX4-KO BALB/c mice with or without 10 µM ATPγs activated with IgG-OVA-coated beads for the indicated times. OVA (green) and Lamp1 (magenta). (**f**) Quantification of OVA fluorescence intensity remaining on beads interacting with cells. (**g**) Quantification of the percentage of Lamp1 MFI in the area enclosed by the bead, normalized by total Lamp1 MFI in the cell. (**h**) Representative images of P2RX4-silenced B cells activated with IgG-CypHer5E(magenta)-coated beads. (**i**) Quantification of the MFI of IgG-CypHer5E-coated beads interacting with cells, normalized by the 1,1 of the average of non-interacting beads. ****P<0,0001 Kolmogorov-Smirnov test n>215 from three independent experiments. (**j**) Antigen presentation (LACK) assay of B cells silenced for P2RX4. B cells were incubated with IgG- or IgM-Lack-coated beads for 2 h or 4 h. Cells were fixed and co-incubated with LMR 7.5 (Lack-specific T cells), for 4 h at 37° C. IL-2 levels were measured by ELISA. (**k**) Peptide control of silenced B cells was used in the antigen presentation assay. Mean amounts of IL-2 are shown for a representative of three independent experiments performed in triplicate. Two-way ANOVA with Sidak’s multiple comparison test. Data consider n ≥ 20 cells pooled from N = 3 independent experiments. Two-way ANOVA with Sidak’s multiple comparison test. P values illustrated with asterisks are ** <0.01, *** <0.001, and **** <0.0001. Error bars are mean ± SEM.

To evaluate the contribution of purinergic receptors in B cell activation, we used suramin, a broad inhibitor of this receptor family^38^. B cells were preincubated with suramin (10-100µM) for 1 h, and antigen extraction was subsequently measured using IgG-OVA-coated beads. Treatment with 50 and 100 µM suramin significantly reduced antigen extraction, suggesting this family of purinergic receptors may be involved in B cell activation (Extended Data Fig. 2b).

Given that our cell line does not express P2RX7^39^ we focused on P2RX4, which is expressed in most mature B cell subtypes, while P2RX3 is limited to pre-B cells. Silencing P2RX4 in IIA1.6 B cells using siRNA resulted in a significant reduction in antigen extraction following stimulation with IgG-OVA-coated beads at 60 and 120 min compared to control cells (Fig. 2b,c,d). To further validate the role of P2RX4 in antigen extraction, we isolated primary naïve B cells from the spleens of P2RX4-KO mice. Cells were incubated with antigen-OVA-coated beads, and the remaining fluorescence intensity of OVA was measured at various time points. Consistent with our previous results, P2RX4-KO cells showed reduced antigen extraction compared to WT cells at early activation times (30 and 60 min) (Fig. 2e,f,g). However, after 120 min, no significant differences were observed between the two groups (Extended Data Fig. 2c), suggesting that P2RX4 is not strictly required for antigen extraction but enhances its efficiency, particularly during the early stages of the response. To directly assess whether the effect of extracellular ATP on antigen extraction by B cells depends on P2RX4, we incubated primary B cells from wild-type (WT) and P2RX4-knockout (P2RX4-KO) mice with 10 µM ATPγS. After 30 min of activation, B cells isolated from P2RX4-KO mice exhibited reduced antigen extraction compared to WT controls, and ATPγS treatment did not rescue this defect. By 60 min, however, this difference was no longer observed (Fig. 2e,f,g). Together, these results suggest that the effect of extracellular ATP on antigen extraction by B cells is mediated, at least in part, through P2RX4.

The reduced antigen extraction observed in the absence of P2RX4 can be mechanistically linked to its role in promoting lysosome fusion and/or exocytosis^40^. Here we show that in P2RX4-deficient cell lines, there is an increased accumulation of lysosomes at the synaptic interface coupled with reduced antigen extraction, suggesting a defect not in lysosome recruitment but in their functional deployment at the membrane (Fig. 2d,g). To formally evaluate whether silencing P2RX4 affects lysosome fusion at the IS, we used CypHer-5E-coupled to antigen-coated beads. This probe has maximal fluorescence at acidic pH, which in this setup correlates with local lysosomal exocytosis at the antigen-bead contact site^36,41^.

Thus, we quantified the fluorescence intensity of CypHer-5E-antigen beads in contact with P2RX4-silenced B cells. We found that beads interacting with P2RX4-silenced B cells showed a significantly lower fluorescent fold-change than controls (Fig. 2h,i), indicating reduced lysosome secretion at the antigen contact site. These findings suggest that the P2RX4 plays a role in lysosomal secretion at the synaptic membrane. We next assessed whether the P2RX4 receptor regulates the ability of B cells to process and present the antigen to CD4+ T cells. To this end, P2RX4-silenced B cells were stimulated with IgG-Lack-coated beads. After 2-4h of incubation with these beads, B cells were fixed and co-incubated for 4 h with CD4+ T cells. IL-2 levels in the supernatant, reflecting T cell activation, show that P2RX4-silenced B cells exhibited a ∼50% reduction in antigen presentation to T cells after 2 and 4h of antigen processing compared to control cells, whereas no differences were observed in peptide presentation (Fig. 2j,k). Together, these results confirm that the P2RX4 regulates B cell activation by promoting the ability of B cells to extract and present immobilized antigens. Additionally, our data suggests that the P2RX4 mediates the effect of extracellular ATP on B cell functions.

### Extracellular ATP regulates actin cytoskeleton dynamics in B cells

B cell activation during antigen recognition involves actin cytoskeleton-dependent spreading responses that increase the contact area with the APC and enhance BCR-antigen encounters^21,42^. Thus, actin cytoskeleton dynamics are crucial for B cell activation^43^, however, the contribution of purinergic signaling to this process remains largely unknown. To examine cytoskeleton rearrangements during B cell activation, cells were seeded onto IgG-coated coverslips and the effects of extracellular ATP on B cell spreading were assessed by staining F-actin with phalloidin. In the presence of 10 µM ATP, B cells exhibited significantly increased cell spreading after 15 (Fig. 3a,b) and 60 min of activation compared to control conditions and after 60 min; however, no differences were observed at 30 min (Extended Data Fig. 3a). As the effect was not linear in time, we suspected that ATP was being degraded, thus we used ATPγS. Cells were pre-incubated with ATPγS for 10 min prior to BCR stimulation, resulting in a similar increase in spreading at 15 min (Fig. 3c,d), consistent with the effects observed with ATP. Notably, at 30 min, ATPγS-treated B cells displayed a reduced spreading area compared to controls, while maintaining a size comparable to that observed at 15 min. After 60 min, control B cells contracted to an area comparable to that of ATPγS-treated cells, which remained stable from 30 min onward (Extended Data Fig. 3b). Overall, our results suggest that ATPγS accelerates the spreading response of B cells. We therefore assessed the contribution of ATP released by B cells in the spreading response by treating the cells with apyrase. We found that 5 U/mL apyrase reduces the B cell spreading response, whereas 1 U/mL does not produce a significant effect compared with control cells (Fig. 3e,f), suggesting that B cells release substantial amounts of ATP to support autocrine activation. We next assessed the impact of P2RX4 depletion on B cell spreading using siRNA. Silencing P2RX4 significantly reduced the spreading area after 30 min of cell activation compared to control siRNA cells (Fig. 3g,h). However, in primary B cells isolated from P2RX4-KO mice, no significant differences in spreading were observed compared to WT cells, suggesting potential compensatory mechanisms in the absence of P2RX4 (Fig. 3i,j)

**Figure 3:**
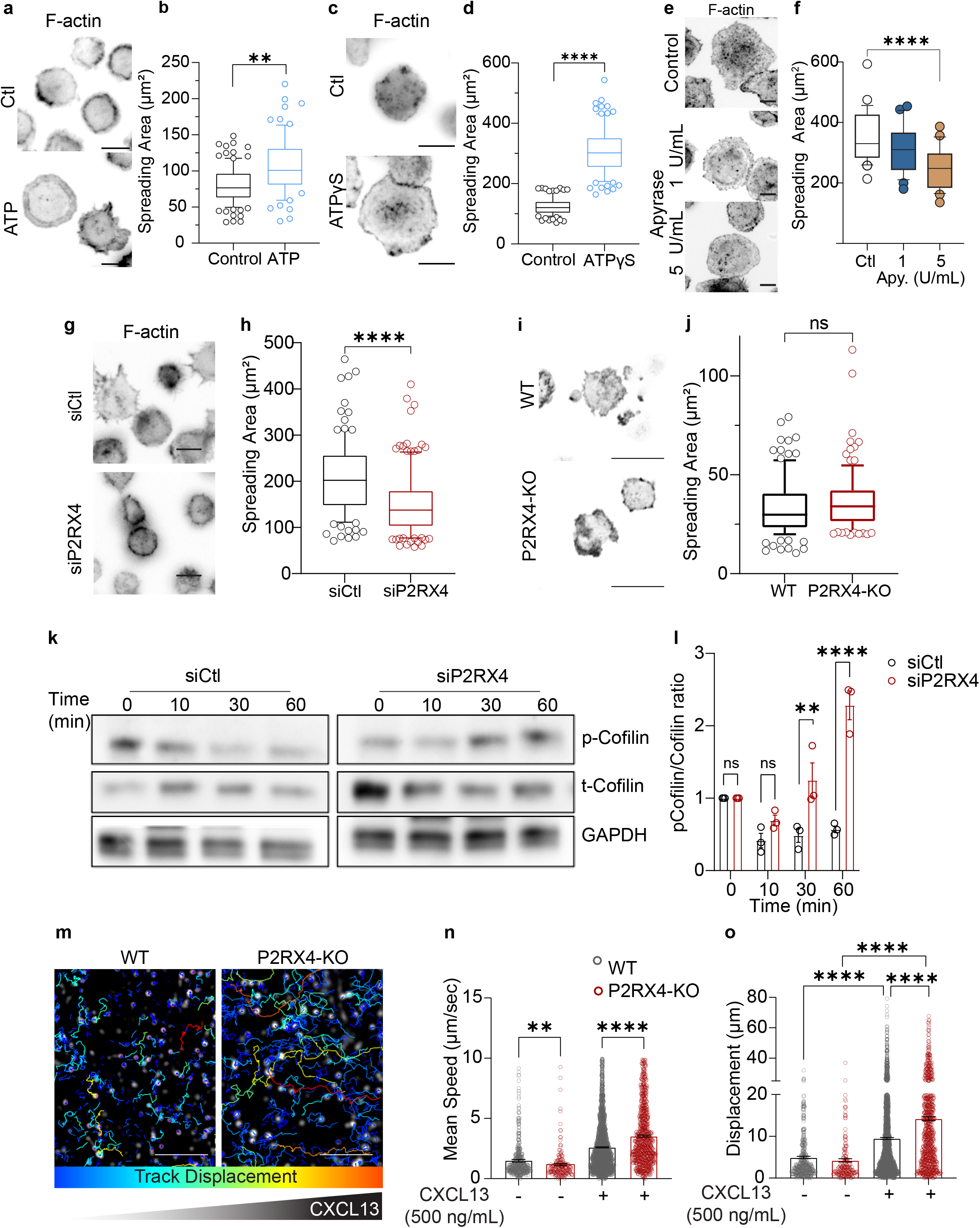
Actin cytoskeleton dynamics in B cells upon stimulation with extracellular ATP. (**a-d**) Representative widefield images of B cells activated on IgG-coated coverslip for 15 min (**a**, **b**) or 60 min (**c**,**d**,**e**) and quantification of cell spreading areas defined by actin staining using phalloidin for B cells treated with: (**a**) ATP (10 µM), (**b**) ATPγS (10 µM), (**c**) 1 and 5 U/mL of apyrase, (**d**) P2RX4 silenced B cells, and (**e**) primary B cells isolated from the spleen of wild-type and P2RX4-KO BALB/c mice. Scale Bar of 10 µm. Data consider n ≥ 20 cells pooled from N = 3 independent experiments. Two-way ANOVA with Sidak’s multiple comparison test. P values illustrated with asterisks are ** <0.01, *** <0.001, and **** <0.0001. Error bars are mean ± SEM. (**f**) Representative western blot to detect expression levels of P2RX4 receptor in silenced and control B cells. GAPDH was used as a loading control. (**g**) Quantification of protein levels from western blot images **P=0,0020 and ****P<0,0001 two-way ANOVA with Šídák’s multiple comparisons test from three independent experiments (**h**) Representative migration tracks of WT and P2RX4-KO BALB/primary B cells in response to CXCL13 in a 3D collagen gel migration assay. Scale bar: 200 µm. (**i**) Left: Quantification of cell displacement before and after chemokine stimulation, analyzed using a multiple-comparisons test. Right: Quantification of the mean cell speed before and after chemokine stimulation

To explore how P2RX4 regulates cytoskeleton-dependent spreading in B cells, we focused on actin regulators that depend on increases in cytosolic calcium as the P2RX4 is a calcium ionotropic channel^44^. For this, we focused on cofilin, an actin-severing protein regulated by calcium signaling through the activation of calcineurin^21,45^. In B cells, cofilin mediates filament severing and depolymerization, driving the cytoskeletal remodeling necessary for BCR signaling and antigen gathering. Cofilin is activated by dephosphorylation, allowing it to remodel cortical actin^46^. Silencing P2RX4 in B cells resulted in sustained phosphorylation of cofilin (inactive form) at 30 and 60 min of activation (Fig. 3k,l), indicating a potential role for P2RX4 in cofilin-dependent actin dynamics during B cell activation.

Actin rearrangements are critical for B cell migration, enabling these cells to scan the lymph node for antigens^47^. Given that silencing P2RX4 affected cofilin activation, we hypothesized that P2RX4 could also influence the migratory behavior of B cells. To test this, we measured the migratory capacity of splenic B cells isolated from P2RX4-KO mice within a 3D collagen matrix that mimics the physiological microenvironment^48^. Upon random migration conditions (i.e., in the absence of a chemoattractant), P2RX4-KO B cells displayed a slight reduction in migration speed compared to wild-type B cells, although their overall displacement was not significantly altered. We then introduced CXCL13, a key chemokine guiding B cell trafficking, using a microfluidic device that generates a stable gradient and promotes directional migration. Strikingly, in the presence of CXCL13, P2RX4-KO B cells exhibited increased displacement and migrated faster than wild-type cells (Fig. 3m,n,o). These findings suggest that P2RX4 plays a role in regulating cytoskeleton-dependent processes, including directional migration and cell spreading responses, both essential for efficient antigen extraction *in vivo*.

### P2RX4 is recruited to the IS of B cells in vamp7-positive vesicles

So far, we have shown that silencing the P2RX4 impairs the ability of B cells to extract immobilized antigens. Next, to investigate the cellular mechanisms involved, we examined the subcellular localization of P2RX4 in B cells during IS formation. To this end, B cells expressing a P2RX4-HA construct were activated with BCR ligand-positive coated beads, followed by staining for the HA-tag to assess the localization of the receptor. In resting conditions, the P2RX4 was localized in vesicular compartments (Fig. 4a and Extended Data Fig. 4a). Upon BCR activation, however, P2RX4 was progressively recruited to the IS. This redistribution was confirmed by an increased polarity index, reflecting its enrichment toward the synapse, as well as by elevated mean fluorescence in the vicinity of the activating bead, indicating accumulation of the receptor at the IS over time (Fig. 4a,b,c and Extended Data Fig. 4a,b,c).

**Figure 4:**
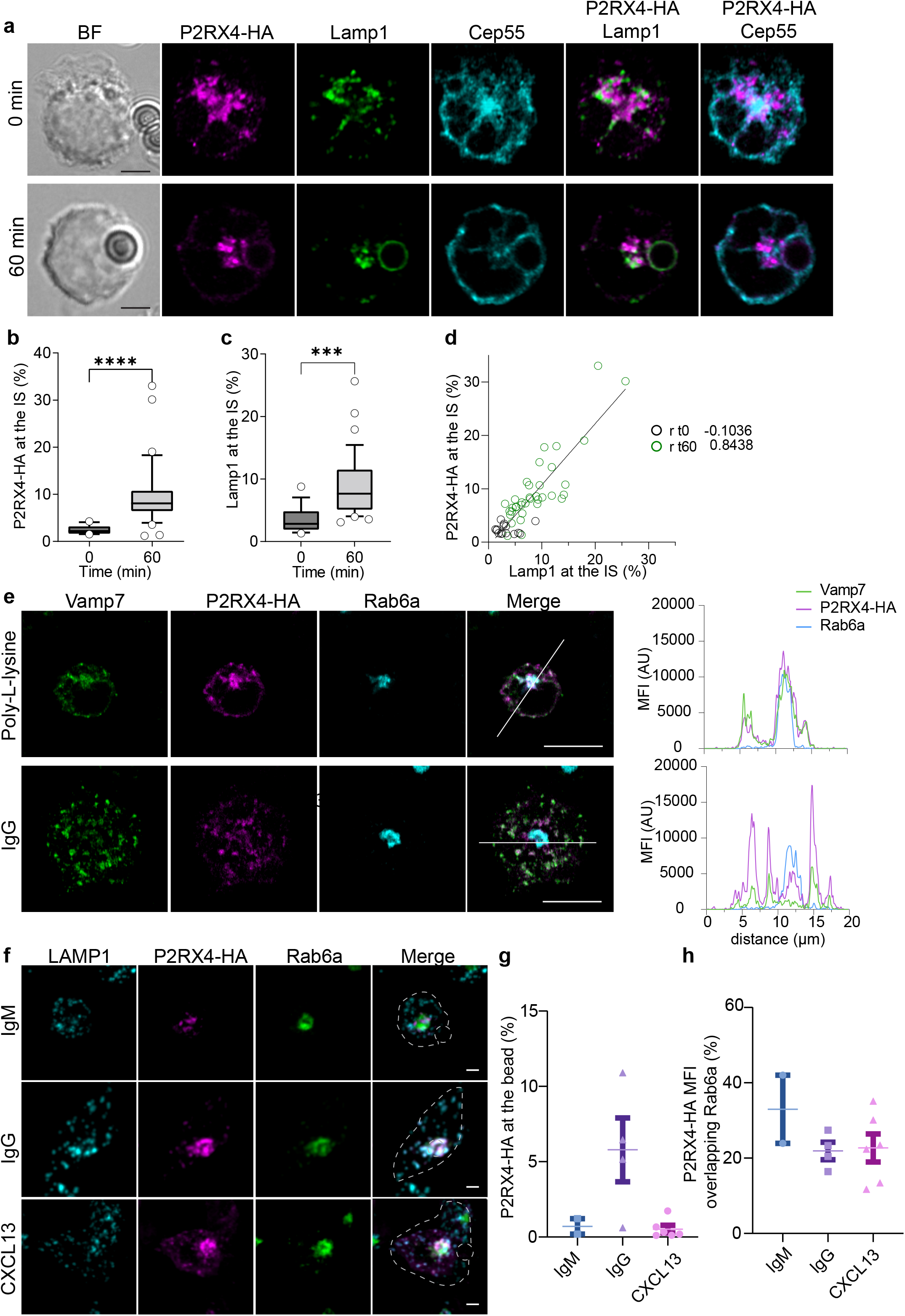
P2RX4 receptor dynamics during B cell activation. (**a-c**) Confocal images of B cells expressing the P2RX4-HA construct (magenta) and co-stained with Lamp1 (lysosomes, green) and Cep55 (microtubules, cyan) stimulated with IgG- or IgM-coated beads for 60 min. Scale bar: 3 µm. Quantification of the percentages of P2RX4-HA MFI (**b**) or Lamp1 (**c**) within the area enclosed by the bead, normalized to the total MFI levels in the cell. 14 cells (time 0) N=2. Kruskal-Wallis test followed by Dunn’s test. (**d**) Spearman correlation (r) of the accumulation of P2RX4-positive vesicles and lysosomes (Lamp1) at the IS of B cells. (**e**) Representative confocal images of B cells expressing the P2RX4-HA construct (magenta) and Vamp7-GFP (green) co-stained with Rab6 (cyan) seeded on glass coverslips containing IgG or poly-L-lysine for 60 min. Right: fluorescence intensity distribution. (**f**) Representative confocal images of B cells expressing the P2RX4-HA construct (magenta) and co-stained with Lamp1 (lysosomes, cyan) and Rab6a (Golgi, green) activated for 60 min with IgG-, IgM- or CXCL13-coated beads. (**g**) Quantification of the percentage of the MFI of P2RX4-HA in the area enclosed by the bead, normalized by the total MFI levels of the cell. (**h**) Quantification of the percentage of the MFI of P2RX4-HA overlapping with the Rab6a compartment, normalized by the total MFI of the cell.

P2RX4 has been previously reported to localize to lysosomal compartments^44,49^. To investigate whether this localization is conserved in B cells, we co-stained the P2RX4-HA with the lysosomal marker Lamp1 and monitored their recruitment to the antigen contact site at different time points following cell activation. Spearmans’ analysis showed an increased correlation between P2RX4-HA and Lamp1 recruitment at the IS upon BCR activation (Fig. 4d). Closer inspection using high-resolution microscopy revealed two types of resting cells: one showing P2RX4-HA localized near the MTOC (stained with Cep55), and another displaying P2RX4-HA associated to the plasma membrane. In both cases, P2RX4-HA partially colocalized with Lamp1 as no significant difference in Pearsons’ and Manders’ coefficient was observed between IgG or IgM stimulated B cells (Extended Data Fig. 4d,e). Upon activation, however, P2RX4 robustly accumulated at the IS, forming a ring-like structure surrounding the antigen-coated bead, closely mirroring the distribution of Lamp1 (Fig. 4a). This pattern is consistent with lysosome docking and fusion events at the synapse as described^36,50^.

To further confirm the presence of P2RX4 at the synaptic membrane, we employed total internal reflection fluorescence (TIRF) microscopy to selectively visualize the synaptic membrane (Extended Data Fig. 4h). We sequentially adjusted the evanescent field depth from 200 nm to 150 nm and finally to 100 nm, thereby progressively restricting visualization to regions closest to the plasma membrane. P2RX4 signal remained clearly detectable across these depths, indicating its presence within the synaptic interface (Extended Data Fig. 4h). This stepwise approach confirmed that P2RX4 reaches the synaptic membrane and, together with our previous observations, supports a functional role for P2RX4 at the IS during B cell activation. To further investigate the distribution of P2RX4 at the IS, B cells were activated on antigen-coated coverslips for different time points (10-60 min). Activation triggered a characteristic spreading response, as evidenced by an increased cell area visualized by F-actin staining. Concomitantly, lysosomes were recruited to the synaptic plane as shown by the staining of Lamp1 (Fig. 4f). Consistent with our previous observations, P2RX4-positive vesicles progressively accumulated at the synaptic interface after 30- and 60-min. Z-axis analysis confirmed this redistribution, revealing a shift of P2RX4 signal from upper cellular regions toward the synaptic membrane over time (Extended Data Fig. 4f,g,h). Additionally, the images show that several P2RX4 positive vesicles localized adjacent to Lamp1 positive vesicles at the center of the IS (Fig. 4f).

Considering that the P2RX4 receptor is rapidly internalized from the plasma membrane to endolysosomal compartments^44^, we next sought to define the vesicular compartments harbouring P2RX4 during B cell activation. For this purpose, we performed triple labelling of resting and activated B cells after 60 min with P2RX4-HA, lysosomes (Lamp1) and early endosomes (Rab5-YFP) and performed colocalization analysis. Notably, in resting conditions imaging analysis showed that P2RX4 colocalized more with early endosomes than with lysosomes (Pearson’s 0.83 and 0.069, respectively). This preferential association was no longer observed following B cell activation (Extended Data Fig. 4i,j). We further examined P2RX4 localization relative to additional trafficking markers, including VAMP7, a vesicular SNARE associated with Lamp1 positive vesicles and required for lysosome fusion at the B cell synapse^50^ and Rab6a, a GTPase localized to the trans-Golgi network^51^. Using confocal Airyscan microscopy, we observed that under non-activating conditions, a pool of P2RX4 overlaps with the Golgi apparatus from where it becomes depleted upon activation, similarly to VAMP7 (Fig. 4e). Following activation, a substantial fraction of VAMP7-positive vesicles also contained P2RX4, although a subset of vesicles remained exclusively P2RX4-positive, indicating the presence of distinct trafficking populations. Overall, these results suggest that P2RX4-positive vesicles relocate from the Golgi apparatus towards the antigen contact site, where they partially coalesce with Vamp7^+^/Lamp1^+^ vesicles.

We next compared the subcellular localization of P2RX4 under BCR-activating, non-activating, and migratory stimulation conditions using CXCL13-coated beads. Consistent with our previous observations, P2RX4 predominantly overlapped with Rab6a-positive compartments under non-activating conditions, whereas BCR activation promoted its recruitment to the bead and decreased its overlap with Rab6a. Under migratory stimulation, no recruitment of P2RX4 to the bead was observed, together with reduced overlap with Rab6a and a more even distribution of P2RX4 in lysosomes (Fig. 4f,g,h). These results suggest that P2RX4 intracellular trafficking varies depending on the stimulus. BCR activation induces P2RX4 recruitment to the synaptic membrane, depleting its localization in Rab6a-positive compartments, whereas CXCL13 stimulation reduces P2RX4 association with Rab6a-positive compartments without promoting its recruitment to the bead, leading instead to a more even distribution within non-clustered lysosomes.

## Discussion

In this study, we show that extracellular ATP enhances key B cell functions, particularly their ability to extract and present immobilized antigens at the IS. Mechanistically, this enhancement is not due to increased lysosomal recruitment but rather to increased enzymatic efficiency, suggesting that ATP modulates the functional output of lysosome-mediated antigen processing. Importantly, we identify a pivotal role for purinergic receptor P2RX4 during this process. We further show that P2RX4 regulates actin dynamics through cofilin-1, thereby influencing B cell spreading and motility. Additionally, P2RX4 displays a dynamic intracellular localization, residing in internal compartments under resting conditions when the cells are motile, and trafficking to the IS upon BCR activation when the cells decrease their motility for efficient antigen presentation.

The effect of ATP in B cell biology has been explored, yet studies report different outcomes. One study described a stimulatory role through increased Ca²⁺/PLC signaling, whereas another showed that ATP, via P2X7, reduces Ca²⁺ signaling and impairs NFAT nuclear translocation^19,20^. The effect of extracellular ATP over B cell activation has been analyzed at both early and late stages using a broad range of concentrations that activate different P2 receptors^19,20^. We observed a non-linear effect of ATP: intermediate concentrations (10–50 μM) enhanced antigen extraction, whereas higher concentrations (100 μM) reduced this capacity, suggesting the existence of a functional threshold and the role of different P2 receptors. Furthermore, non-hydrolyzable ATP elicited a stronger response than hydrolysable ATP, indicating that ATP degradation products, such as adenosine, may differentially modulate B cell activity.

On the other hand, we also show a key role for the release of endogenous ATP. We found that B cells locally release ATP at the site of antigen engagement, supporting the idea of polarized ATP secretion. This is consistent with previous observations showing that tonic ATP secretion increases upon B cell stimulation, suggesting that ATP may act in an autocrine manner to promote activation^52,53^, which is finely tuned with their metabolic state ^54^. The autocrine role of ATP is similar to that observed in T cells, which release ATP upon TCR engagement via Panx1 channels thereby activating P2RX1 and P2RX4^55^. Similarly, dendritic cells release ATP through Panx1 channels to sustain P2X7 and subsequent CaMKII activation, which lead to actin remodeling and fast migration^33^. The autocrine release of ATP in B cells is supported by membrane channels, Cx and Panx channels, which are known to mediate ATP release in immune cells^32,33^. However, this could occur concomitantly with lysosomal-dependent ATP release, which might occur during antigen extraction^36,53^. Collectively, these findings highlight polarized ATP release as a conserved mechanism across immune cells, which helps coordinate their responses more precisely, ensuring effective activation when needed while preventing excessive immune reactions.

A key finding of this work is that P2RX4 trafficking is tightly linked to its function. We show that P2RX4 translocates from intracellular compartments, particularly the trans Golgi network, to the IS upon BCR engagement. This trafficking likely occurs via Lamp1 and Rab5-positive vesicles, and resembles the behavior of the snare protein Vamp7, which regulates lysosomal fusion at the synaptic membrane^50^. We propose a model where lysosomal fusion inserts P2RX4 into the plasma membrane, exposing its ATP-binding domain to the extracellular environment (Fig 5). Given that lysosomal pH inhibits P2RX4 activity, fusion with the plasma membrane or less acidic compartments may relieve this inhibition, enabling receptor activation^44,49^. Once activated, P2RX4 promotes calcium influx, which supports both lysosomal fusion and actin remodeling^56,57^. Functionally, this places P2RX4 as a key regulator of IS formation. In line with this, we evaluated B cell migration in response to the chemokine CXCL13, which is produced within lymph node follicles and guides B cells as they patrol the follicular environment in search of antigen^58^. Under these conditions, we observed that P2RX4-deficient B cells exhibit increased migration in response to CXCL13 but display reduced spreading and antigen extraction triggered by BCR activation. These findings suggest that P2RX4 functions as a molecular switch that restrains chemokine-driven motility while promoting IS formation. In this way, P2RX4 may coordinate the transition between migratory behavior and stable antigen engagement, halting B cells at sites of antigen encounter to facilitate efficient recognition and extraction.

**Figure 5:**
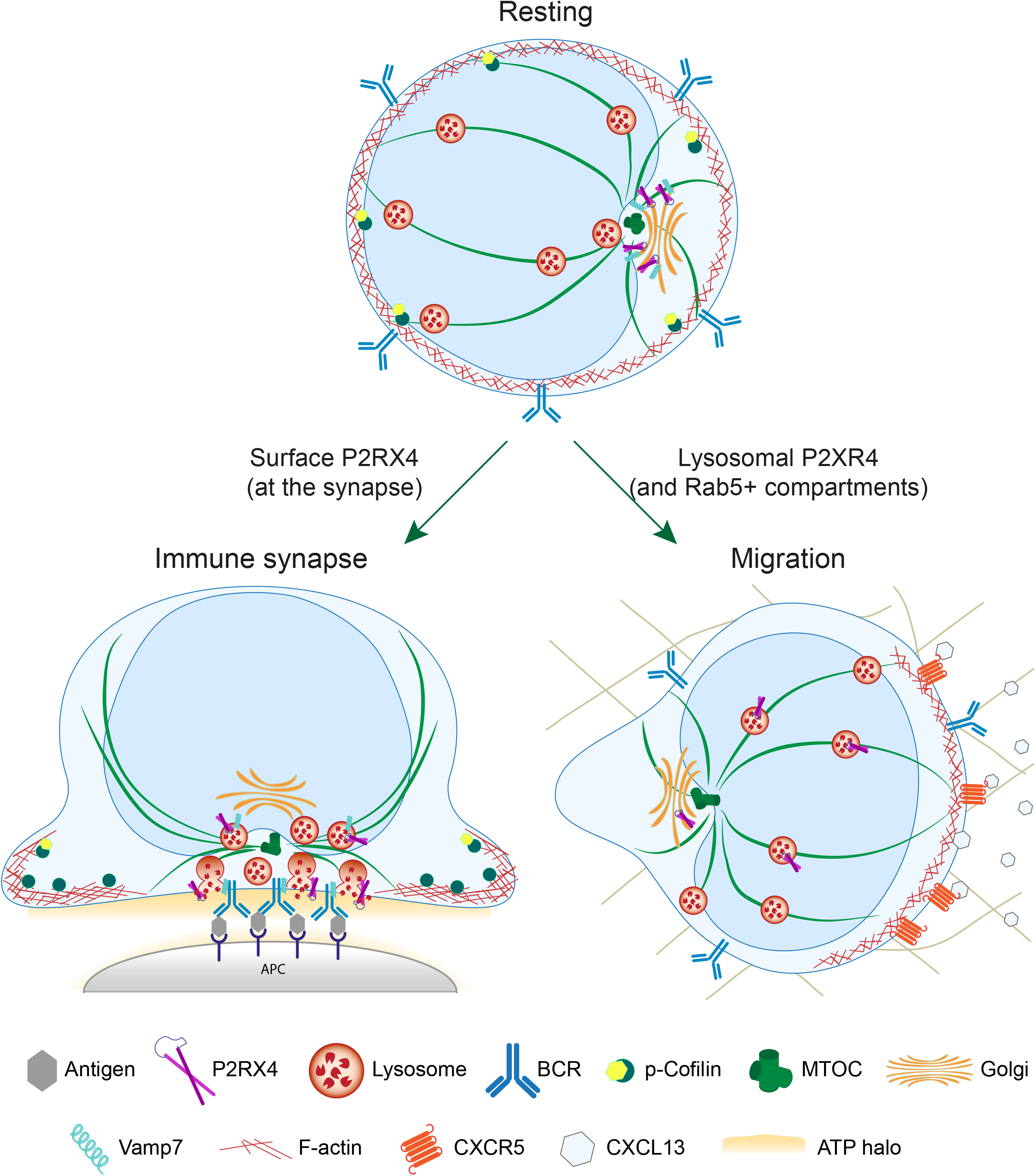
Schematic representation of the function of P2RX4 in B cells. Under resting conditions, P2RX4 resides in the Golgi apparatus. Upon BCR engagement, P2RX4 traffics to the IS in VAMP7-positive vesicles, where it is inserted into the plasma membrane with its ATP-binding domain exposed extracellularly. BCR activation induces local ATP release, enabling activation of synaptic P2RX4. In addition, extracellular ATP derived from non-B cell sources further enhances B cell activation by increasing cell spreading, antigen extraction, and presentation. P2RX4 signaling promotes cofilin activation (dephosphorylation), which regulates actin remodeling for B cell spreading, thereby supporting efficient antigen extraction and presentation on MHC-II. Conversely, P2RX4 restrains B cell migration under chemotactic CXCR5-CXCL13 axis, suggesting that P2RX4 functions as a molecular switch coordinating the balance between immune synapse formation and cell migration.

ATP-induced cytoskeleton remodeling was P2RX4-dependent, which likely relies on local calcium signals that might lead to actin remodeling at the IS^56^. Elevated intracellular calcium levels activate calcineurin, which subsequently enhances the activity of the phosphatase Slingshot (SSH), resulting in cofilin dephosphorylation and activation^45^, as observed in our model. Sustained levels of phosphorylated cofilin impair actin turnover, resulting in excessive F-actin accumulation, reduced cytoskeletal flexibility, and defective BCR microcluster formation, ultimately limiting the ability of B cells to efficiently extract immobilized antigens^59^. Importantly, actin remodeling is also required for lysosome docking and fusion^60^, indicating that P2RX4 is most likely involved in multiple steps associated with antigen extraction. Similar roles for P2RX4 have been described in other immune cells, where it regulates lysosomal dynamics, vesicle fusion, and phagocytosis^49^. Thus, P2RX4 may represent a conserved regulator of membrane trafficking and cytoskeletal organization across leukocytes.

Other purinergic receptors may contribute to ATP-mediated regulation of B cell functions. B cells express low P2RX7 compared to T cells^61^ , and our cellular model lacks P2RX7 expression^39^, allowing us to specifically investigate the role of P2RX4. While P2RX1 could also participate in modulating B cell responses, as shown during T cell activation^55^, its low expression or absence in B cells makes it unlikely. P2RY11 metabotropic receptor regulates leukocyte migration, cytokine production, and responses to inflammatory signals^62,63^. More broadly, P2Y receptors are known to control processes such as chemotaxis, phagocytosis, and immune cell communication through G protein-coupled signaling pathways^64^. These receptors may act in concert with ionotropic P2X receptors to integrate extracellular nucleotide signals, thereby fine-tuning immune responses. Future studies should address the relative contribution and interplay of P2X and P2Y receptors in B cells.

Extended Figure 1.

(**a**) Representative confocal images of B cells expressing the ATP sensor (pm-iATPsnFR1.1) seeded on glass coverslips containing IgG or poly-L-lysine after 60 min.

(**b**) Z-scan profile of the MFI of the ATP sensor in B cells stimulated with IgG or poly-L-lysine for 60 min.

(**c**) MFI of the ATP sensor at the IS per condition.

Quantification of OVA fluorescence intensity remaining on beads interacting with cells treated with: (**d**) Cbx, (**e**) ^10^Panx1 and (**f**) La^3+^, respectively. Values were normalized by fluorescence at time zero.

Representative widefield images of B cells in the presence of: (**g**) 1 and 5 U/mL of apyrase, (**i**) 0 and 10 µM of ATP or (**r**) Primary B in the presence of 10, 50, and 100 µM ATPγs activated with IgG-OVA-coated beads for the indicated times. OVA (green) and Lamp1 (magenta). Scale bars are 3 μm.

(**h**) and (**j**) Quantification of OVA fluorescence intensity remaining on beads interacting with B cells under different conditions. Values were normalized by fluorescence at time zero.

(**k,o** and **t**) Quantification of the percentage of the MFI of Lamp1 in the area enclosed by the bead normalized by MFI in the cell.

(**l**) Gene heatmap expression from Immgen.org database for enzymes Entpd1 (CD39) and Nt5e (CD73) in B-cell populations.

(**m**) Quantification of the percentage of B cells expressing the ectonucleotidases CD39 and CD73 on the cell surface of resting and activated B cells, and total levels in permeabilized B cells.

(**p**) Quantification of the MFI of Magic red at the synapse plane.

(q) Representative images of B cells stained with Magic Red in the presence of 10, 50, and 100 µM ATPγs. Cells were seeded onto IgG-coated coverslips for 60 min. Kruskal -Wallis test.

Data shown in (**d, e, f, h, j, k, o and t**) consider n ≥ 30 cells pooled from N = 3 independent experiments. Two-way ANOVA with Sidak’s multiple comparison test. P values illustrated with asterisks are ** <0.01, *** <0.001, and **** <0.0001. Error bars are mean ± SEM. *t* test. P values illustrated with asterisks are ** <0.01, *** <0.001, and **** <0.0001. Error bars are mean ± SEM.

Extended Figure 2.

(**a**) Gene heatmap expression of P2X receptor family in immune cell populations (from Immgen.org database).

(**b**) Quantification of OVA fluorescence intensity remaining on beads interacting with B cells treated with 10 and 50 µM of Suramin (a non-selective P2X inhibitor) for the indicated times. Values were normalized by fluorescence at time zero.

(**c**) Representative widefield images of primary B cells, WT or P2RX4-KO (C57BL/6) activated with IgG-OVA-coated beads for the indicated times. OVA (green) and Lamp1 (magenta).

(**d**) Quantification of OVA fluorescence intensity remaining on beads interacting with wild type and P2RX4-KO B cells.

(**e**) Quantification of the percentage of the MFI of Lamp1 in the area enclosed by the bead normalized by MFI in the cell.

Data shown in (**b, d and e**) consider n ≥ 30 cells pooled from N = 3 independent experiments. Two-way ANOVA with Sidak’s multiple comparison test. P values illustrated with asterisks are ** <0.01, *** <0.001, and **** <0.0001. Error bars are mean ± SEM. *t* test. P values illustrated with asterisks are ** <0.01, *** <0.001, and **** <0.0001. Error bars are mean ± SEM.

Extended Figure 3.

Widefield images of B cells activated on antigen-coated coverslips for the indicated times with its respective quantification of cell spreading area defined by actin staining using phalloidin for: (**a**) IIA1.6 B cells treated with ATP (10 µM), and (**b**) B cells treated with ATPγS(10 µM).

Data shown consider n ≥ 30 cells pooled from N = 3 independent experiments. Two-way ANOVA with Sidak’s multiple comparison test. P values illustrated with asterisks are ** <0.01, *** <0.001, and **** <0.0001. Error bars are mean ± SEM.

Extended Figure 4.

(**a**) Widefield images of B cells expressing the P2RX4-HA construct (green) and co-stained with Lamp1 (lysosomes, magenta) activated with IgG-coated for the indicated times.

(**b**) Quantification of the polarity index calculated for P2RX4 receptor.

(**c**) Quantification of the percentage of the MFI of P2RX4-HA in the area enclosed by the bead normalized by the total MFI in the cell.

(**d**) Confocal images of B cells expressing the P2RX4-HA construct (green) and co-stained with Lamp1 (lysosomes, magenta) stimulated with beads coated with BCR-positive and - negative ligand for 60 min.

(**e**) Left: quantification of Pearsons’ coefficient for P2RX4 receptor and Lamp1. Right: quantification of Manders’ coefficient for P2RX4 receptor overlap with Lamp1.

(**f**) Confocal images of B cells expressing the P2RX4-HA receptor construct (green), activated on antigen-coated coverslips for the indicated times. The planes of cells in contact with the glass are shown. P2RX4 (green) and Lamp1 (magenta). Scale bar: 10 µm.

(**g**) Top: visualization of the P2RX4 receptor distribution at different cell planes using the Temporal-color code plugging (cold colors indicate contact with the glass, while warm colors are upper cell planes). Bottom: quantification of P2RX4 receptor distribution in the z-planes of the cell. The grey rectangle represents the region of the IS close to the antigen.

(**h**) Representative images of P2RX4 receptor at the IS plane by total internal reflection microscopy after 60 min of activation on antigen-coated coverslips. The image shows the same cells with decreasing laser depth: 200, 150 and 100 nm. Scale bar: 10 µm.

(**i**) Confocal images of B cells expressing the P2RX4-HA receptor construct (magenta) and the early endosome marker Rab5-GFP, activated on antigen-coated coverslips for the indicated times. The planes of cells in contact with the glass are shown. P2RX4 (magenta), Lamp1 (cyan), Rab5 (green), and phalloidin (grey). Scale bar: 10 µm.

(**j**) Quantification of Manders’ coefficient for P2RX4 receptor overlapping with Lamp1 and Rab5.

Data shown consider n ≥ 30 cells pooled from N = 3 independent experiments. Two-way ANOVA with Sidak’s multiple comparison test. P values illustrated with asterisks are ** <0.01, *** <0.001, and **** <0.0001. Error bars are mean ± SEM. *t* test.

## MATERIALS and METHODS

### Mice, cells, and cell culture

The mouse IgG+ B-lymphoma cell line IIA1.6 ^28^ and the LMR7.5 T cell hybridoma that recognizes I-Ad-Lack156–173 complexes ^65,66^ obtained from A.-M. Lennon-Dumenil. The cells were cultured in CLICK medium (RPMI 1640 with Glutamax supplemented with 10% heat-inactivated FBS, 0.1% 2-mercaptoethanol, 100 U/ml penicillin, 100 μg/ml streptomycin, and 1 mM sodium pyruvate). All cell culture products were purchased from Life Technologies. Spleen B cells were isolated from WT and P2RX4 KO C57BL/6 and BALB/c mice using a magnetic-activated cell sorting B cell isolation kit (Miltenyi) with BMACS columns (depletion of LBs expressing CD43: activated LBs, plasma cells and B-1a CD5+ cells; and all non-LB immune cells), according to the manufacturer’s instructions. P2RX4 KO mice were obtained in collaboration with Prof. Jörg Heeren from the SFB1328, and organs were collected under protocol ORG 1062 granted to Prof. Pablo J. Sáez by the Animal Welfare Officers of the University Medical Center Hamburg-Eppendorf (UKE).

### Reagents

Adenosine 5′-Triphosphate, Disodium Salt (ATP, Sigma-aldrich, Cat#1191, dissolved in water), ATPγS tetralithium salt (ATPγs, Tocris Bioscience, Cat#4080 dissolved in water), suramin sodium salt (Sigma-aldrich, Cat#S2671, dissolved in water), apyrase from potatoes (Sigma-Aldrich #A6535, dissolved in water), ^10^Panx (Bio-techne (rndsystems) #3348/1 dissolved in), Lanthanum(III) chloride heptahydrate (Sigma-Aldrich #262072, dissolved in water), carbenoxolone disodium salt (Cbx, Sigma-Aldrich #C4790, dissolved in water).

### Antibodies and fluorescent probes

The following primary antibodies are used for immunofluorescence: rat anti-Lamp1 (BD Pharmigen #553792), rabbit anti-CEP55 (microtubule marker #ab170414), anti-HA (chicken AB3254 or rabbit #ab137838), rabbit serum anti-OVA (sigma Aldrich #C6534) and rabbit anti-P2RX4 (Alomone labs #APR-024). The following secondary antibodies (1:500) from Jackson Immunoresearch were used: donkey anti-rat F(ab′)2 antibodies conjugated to AlexaFluor488 (#712-546-150), to Cy3 (#712-166-153) and AlexaFluor647 (#712-606-153); together with donkey anti-rabbit F(ab′)2 conjugated to AlexaFluor488 (#711-546-152), to Cy3 (#711-166-152), to AlexaFluor647 (#711-606-152). Donkey anti-chicken F(ab′)2 antibodies conjugated to AlexaFluor488 (#703-546-155) and to Cy3 (#703-165-155).

The antibodies used for western blotting were rabbit: anti-cofilin (1:1000), anti-phospho-cofilin (1:1000), and anti-GAPDH (1:10000); followed by secondary antibodies donkey anti-rabbit coupled to radish peroxidase (1/5000).

The probes used were mitotracker (Invitrogen, #D3922), TMRE (Abcam, #ab113852), ATP red1 (Biozol(OCI), #HY-U00451), PK mito deep red (Spirochrome, #055_23.01), and the pH-sensitive fluorescent probe CypHer5E mono NHS Ester to assess acidification of the synaptic space (Cytiva, #PA15401).

The coupled-antibodies used for flow cytometry were CD73-APC clone TY/11.8 (Biolegend, Cat#127210), CD73-PECy7 clone TY/11.8 (Biolegend, Cat#127224), CD39-PE clone 24DMS1(eBiosciences, Cat#120391) and CD39-PE Texas red clone DUHA59 (Biolegend, Cat#143812)

### Cell electroporation

P2RX4-HA (to assess cellular localization of the receptor, kindly provided by Nelson Barrera), Vamp7-GFP (to assess Vamp7-positive vesicles recruited to the IS), pm-iATPsensorFr1.1 (for visualization of extracellular ATP, addgene #102549). Plasmids were used with Nucleofector R T16 (Lonza, Gaithersburg, MD, United States) to electroporate 1 million IIA1.6 B cells with 1 µg of plasmid. Cells were cultured for 18 h before functional analysis. For P2RX4 receptor silencing, a SMARTPool ON-TARGETplus (DHarmacon #L-048009-00-0005) was used at 300 nM concentration with 2 million IIA1.6 cells 48 h before functional assays. We used a scrambled siRNA (Qiagen) as a control at 10 nM.

### Flow cytometry

B cells were resuspended in PBS + 2% FCS and incubated for 30 min with antibodies against CD39 and CD73 (see antibodies section for antibody details) for surface detection. Under permeabilizing conditions, B cells were resuspended in binding solution (paraformaldehyde 4% in water) and incubated for 30 min at 4 °C. Cells were then washed and resuspended in PBS + 2% FCS for FACS analysis. Analysis of the data obtained from FACS was performed using FlowJoV10 software.

### Surface preparation with BCR +/- ligand

To coat surfaces, an F(ab)2 fragment of the anti-mouse IgG antibody (Biozol (OCI), #JIM-115-006-146) that activates the cell (IgG) and poly-L-lysine was used as a negative control. To coat coverslips, a 100 µg/mL solution of IgG is prepared in PBS 1X, and 45 µL of this solution is added to a parafilm, where the coverslips are deposited. They are incubated overnight at 4°C to bind the ligand by passive absorption and then washed three times with PBS 1X.

To prepare beads with immobilized antigen, 100 µl of Polybead Amino 3 micron (2.5% solids-latex) is used and activated for 4 h with 8% glutaraldehyde in PBS 1X, protected from light at room temperature (RT) in motion. Then, they are washed three times with PBS1X and incubated with 100 µg/mL IgG or an F(ab)2 fragment of the anti-mouse IgM antibody (Biozol (OCI), #115-006-075) with OVA at 100 µg/mL for antigen extraction assays or *Leishmania major* antigen, Lack (Leishmania homolog of mammalian RACKs, the receptors for activated C kinase) 100 µg/mL for antigen presentation assays, overnight at 4°C with agitation. Nonspecific sites are blocked with 500 µL of filtered BSA 10 µg/µL for 1 h at 4°C with agitation. The beads are washed three times with PBS 1X and resuspended in a final volume of 40 µL PBS 1X.

### B cell stimulation and immunofluorescence

B cells are stimulated with antigen-coated beads in a 1:1 ratio (cells: beads) and seeded on poly-L-lysine coated coverslips for different times at 37 °C. B cells are fixed with 4% paraformaldehyde for 10 min at RT, then incubated in a blocking medium (0.3M glycine plus 2% BSA in PBS1X) for 10 min. Intracellular labelling is performed by incubating the primary antibodies in a permeabilisation buffer (2%BSA with 0.25% of saponin in PBS1X) overnight at 4°C. The coverslips are then washed three times with PBS1X for subsequent incubation with secondary antibodies for one h at RT. Finally, the coverslips are washed three times with PBS1X and mounted on slides using 5 µL of Fluoromount G (Electron Microscopy Sciences #17984-25), which are then left to dry at RT ON.

### Antigen extraction assay

IIA1.6 cells were incubated in a 1:1 ratio with IgG plus protein OVA-coated beads (IgG-OVA-coated beads) and then plated on poly-L-lysine coverglass at 37 °C, fixed and stained for OVA. The OVA remaining on the beads was calculated by establishing a 3µm diameter circle around beads in contact with cells and measuring fluorescence on three-dimensional (3D) projections obtained from the sum of each plane. The percentage of antigen OVA extracted was estimated by the percentage of fluorescence intensity lost by the beads at different time points.

### Antigen presentation assay

10^5^ B-cells were stimulated with IgG- or IgM-coated beads with or without protein Lack (antigen derived from Leishmania Major) for 3 and 4 h at 37 °C. Cells were washed with cold PBS 1X, fixed with cold 0.01% glutaraldehyde for 1 min, and the remaining active sites were inactivated with 100 mM glycine in PBS 1X. The fixed B cells were incubated for 4 h at 37 °C with 1.5*10^5^ LMR7.5 T cells. For peptide control, increasing concentrations of peptide Lack (156-173) were added to cells stimulated with IgG-coated beads. Supernatants were collected, and the IL-2 levels were measured with the BD optEIA Mouse IL-2 ELISA kit (#555148) according to the manufacturer’s instructions.

### B cell migration in collagen gels

Collagen experiments were performed as previously described ^48^. Briefly, primary B cells isolated from the spleen were mixed at 4°C with rat tail collagen type I (Corning) at 3 mg/mL at basic pH and loaded in the custom-made polydimethylsiloxane (PDMS) chamber. The sample was incubated at 37°C for 30 min to allow gel polymerisation. Then, 2 mL of CLICK was added to the dish. Cells were allowed to migrate in a random manner for 1 h, and then 500 ng/mL CXCL13 (R&D Systems) was added to the dish, generating a chemokine gradient that triggered directional B cell migration. Cells were imaged overnight with a DMi8 inverted microscope (Leica) at 37 °C with a 5% CO_2_ atmosphere and a × 10 dry objective (NA 0.40 phase). The resulting movies were processed with average subtraction, mean filter, and Gaussian Blur filter to obtain cells as white round objects on a dark background. Tracking was performed with the *Trackmate plugin (Fiji)* in the first 400 μm from the border of the chamber, where the gradient is stable. Tracks of objects moving < 10 μm in length or lasting < 10 min were removed from the analysis to avoid artefacts.

### Western Blot

B cells were lysed at 4°C in radioimmunoprecipitation assay buffer supplemented with protease inhibitor cocktail (Roche). Supernatants were collected, loaded onto gels, and transferred onto polyvinylidene fluoride membrane (Trans-Blot Semi-Dry Transfer Cell; Bio-Rad). Membranes were blocked in 5% non-fat dry milk resuspended in TBS + 0.05% Tween-20 and incubated overnight at 4°C with primary antibodies, followed by a 60 min incubation with secondary antibodies. Western blots were developed with Westar Supernova substrate (Cyanagen), and chemiluminescence was detected using the G:BOX iChemi (Syngene).

### Image acquisition

Widefield images were acquired using a Nikon Eclipse Ti-2 inverted microscope with a 60X/1.25NA oil immersion objective and an iXon Ultra EMCCD camera (Andor Technology). Image acquisition was performed using NIS-Elements Advanced Research software (Nikon Instruments). Images with z-stacks were obtained with a 0.5 μm difference between slices. Confocal images were acquired using either a Zeiss LSM-880 microscope with Airyscan detection (Carl Zeiss Microscopy GmbH), using a 63X/1.4 NA oil-immersion objective, or a Visitron SD-TIRF spinning disk confocal microscope system (Visitron Systems GmbH, Puchheim, Germany) equipped with a 63×/1.49 NA oil-immersion objective. Z-stacks acquired on the LSM-880 were collected using an axial step size of 0.2 μm and a pixel size of 0.07 μm x 0.07 μm and processed in ZEN Black software with standard Airyscan deconvolution settings. Visitron imaging was controlled using VisiView v4 software. Image processing is performed using the Fiji program, based on ImageJ^67^.

### Total internal reflection fluorescence microscopy

Images were acquired in Nikon Ti2Eclipse inverted microscope with a 100x/1.50 NA oil immersion lens and an iXON Ultra EMCCD camera at 37°C. IIA1.6 cells expressing the P2RX4-HA were plated on antigen-coated glass chambers (NuncTM Lab-TekTM II). The evanescent field depth was tuned by modulating the incident angle (θ) of the laser line, allowing us to generate three distinct optical sections of the sample (penetration depths 200, 150 and 100 nm).

### Analysis of acquired images

The single-cell images in the figures were cropped from large fields, and their contrast and brightness were adjusted manually.

### Non-discrete organelle polarity index

The indexes were calculated using a lab-generated macro for ImageJ^68^. Briefly, the area of the cell and latex bead (3 µm in diameter) is manually selected, from which the cell and bead center coordinates (BeadC and CellC, respectively) are extracted. Then, on the Z-projections of the images, the program determines the coordinates representing the mark’s center of mass (MarkC). Then, the program measures the distance between MarkC and CellC (b), the distance between BeadC-CellC (a), and the angle described by MarkC-CellC-BeadC (angle α). The polarity index is obtained from the following formula:

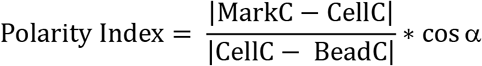

Polarity indexes are observed from a value of -1 (anti-polarized) to 1 (fully polarized).

### Percentage of organelle accumulation at the IS

Organelle recruitment to the IS in bead assays was quantified by dividing the fluorescence at the bead by the total cells’ fluorescence and then multiplied by a factor of 100 to get percentage values.

### Measuring organelle distribution in Z planes

This analysis determines the general fluorescence intensity distribution across the Z-slice of cells seeded onto coverslips (contact slice considered the IS). This measurement assesses the organelle’s proximity to the IS. For this, we first determine the contact slice and then the plane corresponding to the upper limit of the cell, obtaining the MFI of the mark in each z-slice.

## Statistical analysis

Statistical analysis was performed with GraphPad Prism v9 (GraphPad Software; San Diego, CA). The Shapiro-Wilk test was used to contrast the normality of the data set, and then the corresponding analytical test (parametric or nonparametric) was used. P values were calculated using different tests, as indicated in the figure legends.

## Data availability

All data supporting the findings of this study are available within the article and its supplementary information files. Additional materials are available from the corresponding author upon reasonable request.

## ACKNOWLEDGMENTS

We thank N. Barrera for providing the P2RX4-HA plasmid. We thank the Advanced Microscopy Facility at Pontificia Universidad Católica de Chile, especially Nicole Salgado and Fernanda Gárate, for their support with image acquisition. We also thank the UKE Microscopy Imaging Facility (DFG Research Infrastructure Portal: RI_00489), especially Virgilio F., for his support with image acquisition.

This work was supported by a Ph.D. fellowship from the Agencia Nacional de Investigación y Desarrollo #21210344 to M. Alamo Rollandi, a research grant from Fondo Nacional de Desarrollo Científico y Tecnológico #1221128 to M.I. Yuseff, Deutsche Forschungsgemeinschaft project ID 335447717–SFB13228–Project A20 and Human Frontier Science Program grant RGP0032/2022 to P.J. Sáez. We also thank Felipe Del Valle Batalla from the M.I. Yuseff lab for his useful advice concerning experimental procedures and ideas for new approaches.

## Author contributions

**M. Alamo Rollandi:** conceptualization, funding acquisition, data curation, formal analysis, investigation, methodology, software, supervision, validation, visualization, and writing—original draft, review, and editing. **J.P. Bozo:** formal analysis, investigation, methodology, visualization, review, and editing. **I. Riobó:** formal analysis, investigation, methodology, visualization, review, and editing. **T. López-López:** investigation, methodology, review, and editing. **J. Kux:** investigation, resources, review, and editing. **M.Y. Jaeckstein:** resources, review, and editing. **B. Rissiek:** resources, review, and editing. **J. Heeren:** resources, review, and editing. **D. Sauma:** conceptualization, funding acquisition, resources, and supervision. **P. J. Sáez:** conceptualization, funding acquisition, project administration, resources, supervision, visualization, and writing—original draft, review, and editing. **M. I. Yuseff:** conceptualization, funding acquisition, project administration, resources, supervision, visualization, and writing—original draft, review, and editing.

## Disclosures

The authors declare no competing financial interests.

## Notes

### Competing Interest Statement

The authors have declared no competing interest.

