## Supplementary figures and images for "Coupling lysosomal polarity and purinergic signaling regulates B cell activation"

### Extended data

#### Extended Data Figure 1

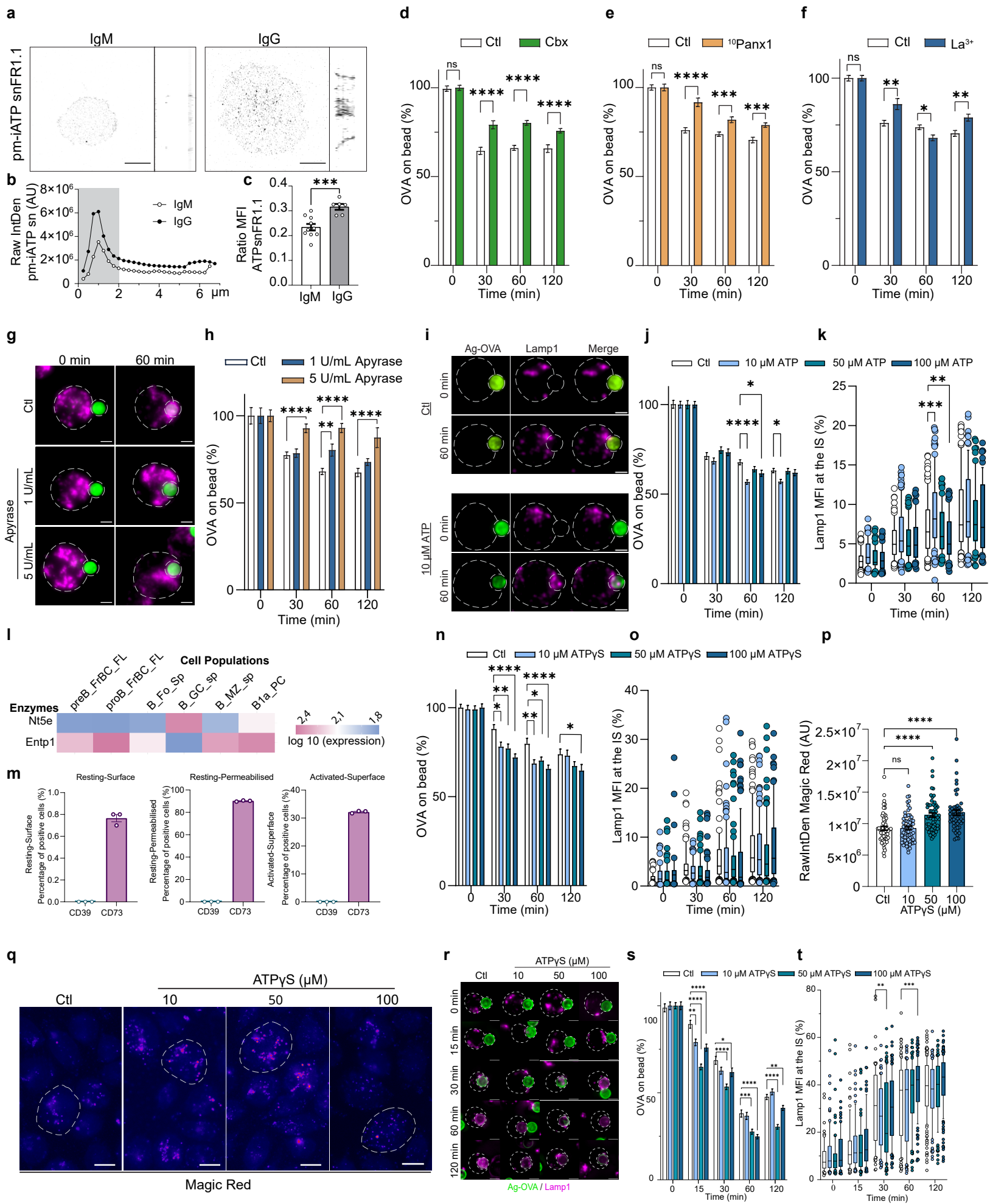

Extended Data Figure 2

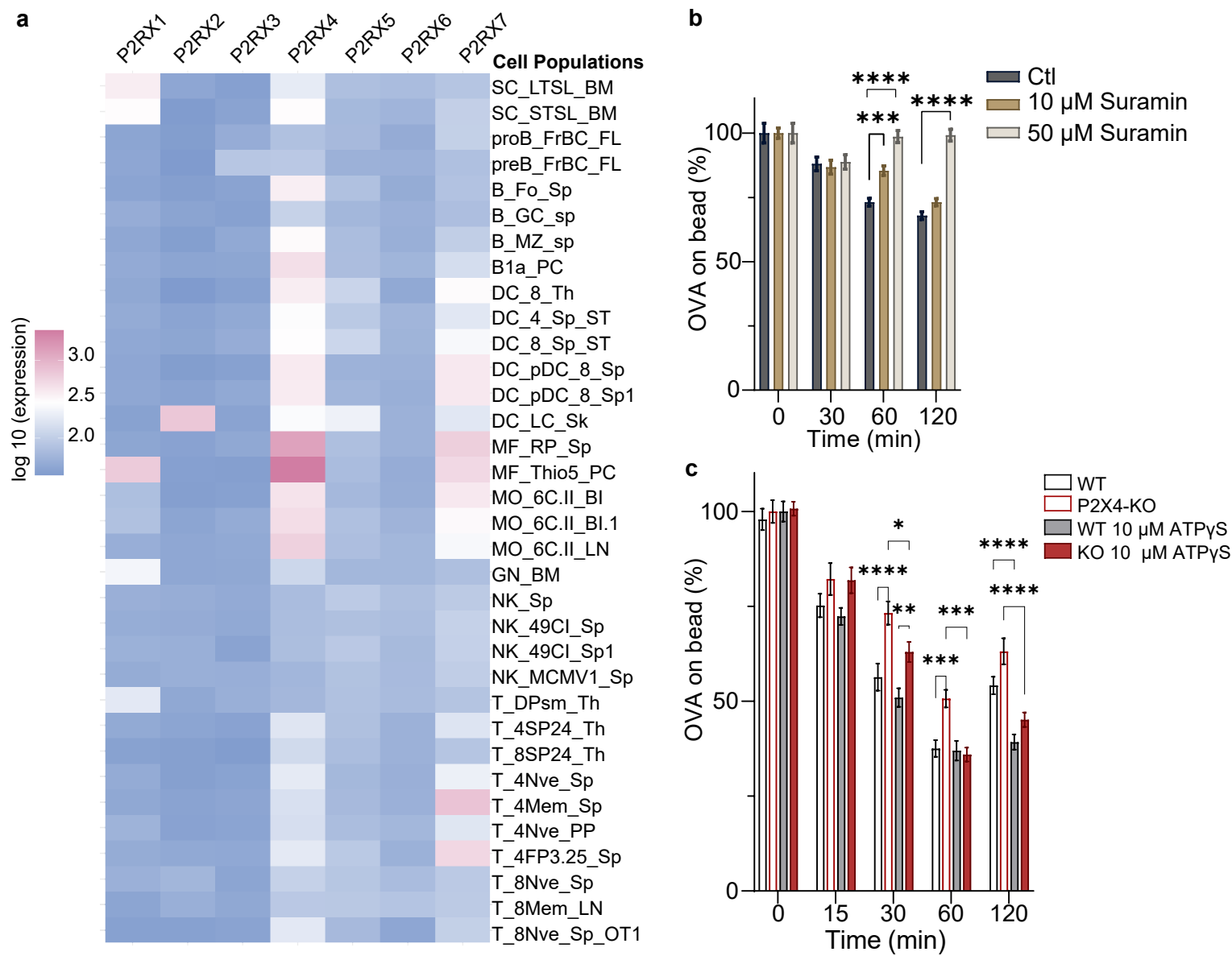

**Extended Data Figure 3**

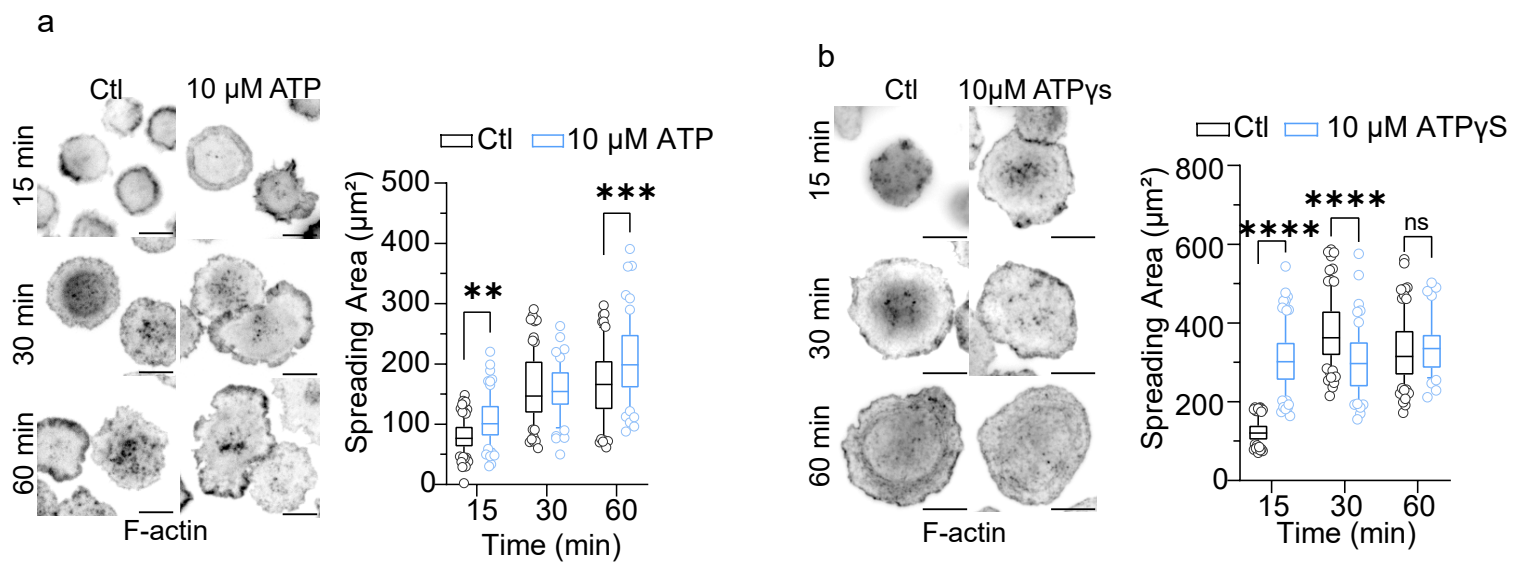

#### Extended Data Figure 4

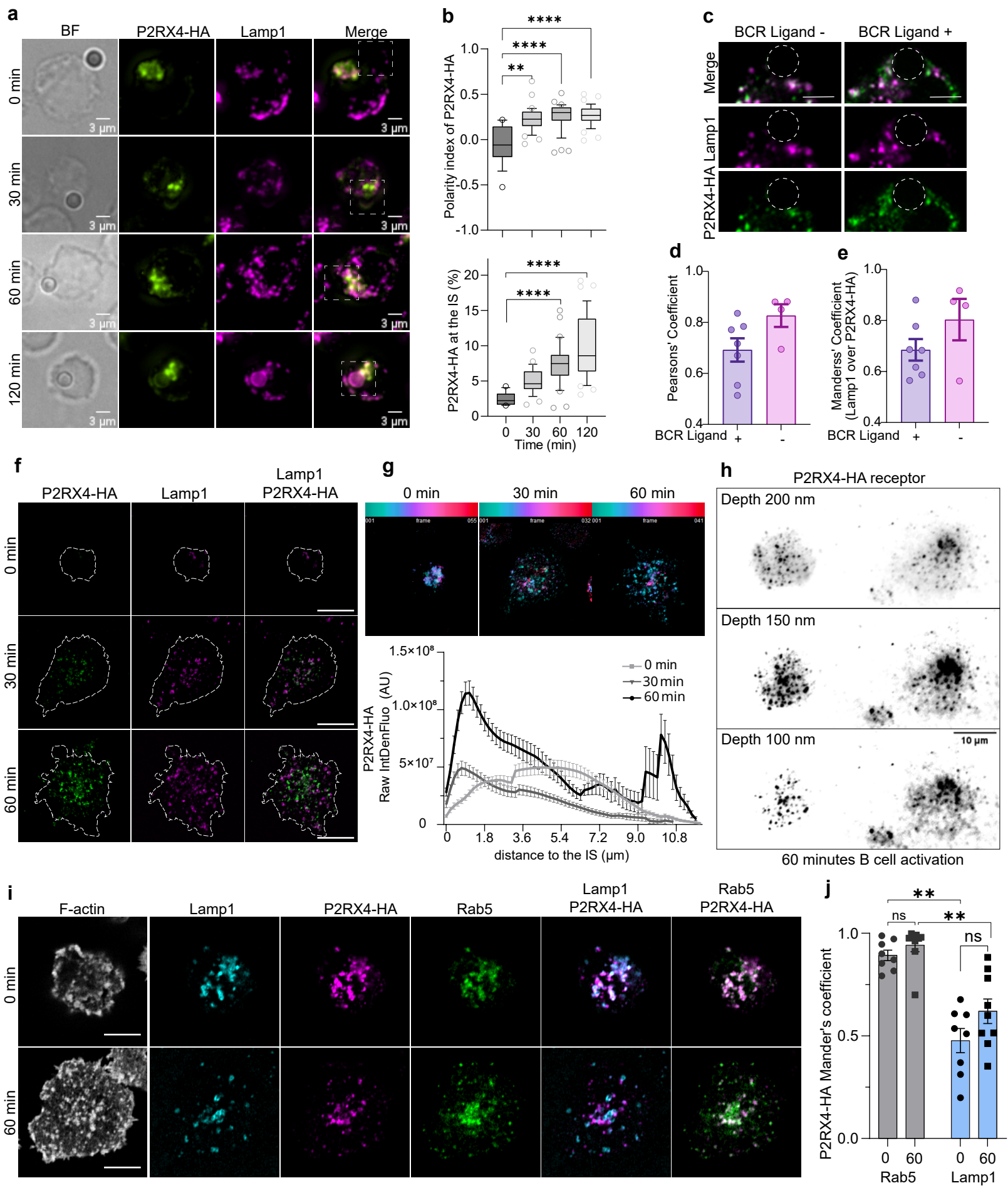
